# Synthesis of extracellular stable gold nanoparticles by *Cupriavidus metallidurans* CH34 cells (revision)

**DOI:** 10.1101/139949

**Authors:** Francisco Montero-Silva

## Abstract

The biogenic synthesis of metallic nanoparticles is of increasing interest. In this report, synthesis of gold nanoparticles by the model heavy metal-resistant strain *Cupriavidus metallidurans* CH34 and *Escherichia coli* strain MG1655 was studied. For the synthesis of AuNPs, bacterial cells and secretomes were incubated with Au(III) ions, revealing that only CH34 cells can produce dispersions of AuNPs. Comparative bioinformatic analysis of proteomes from both strains showed potential CH34 proteins that may be electron donors during reduction of extracellular Au(III) ions and for the biosynthesis of gold nuggets in nature. Powder X-ray diffraction demonstrated that biogenic AuNPs are composed of face-centered cubic gold with a crystallinity biased towards {111} planes. Transmission electron microscopy images showed that AuNPs morphology was dominated by triangular and decahedral nanostructures. EDX and FT-IR spectra showed the presence of sulfur and vibrations associated to the biogenic AuNPs. Based on these results, and analyses of previous genomic and proteomic data, a mechanism for extracellular gold reduction and synthesis of AuNPs by strain CH34 is proposed. Average AuNPs diameter was obtained by nanoparticle tracking analysis, dynamic light scattering and analysis of electron microscopy images. DLS studies showed that biogenic AuNPs colloids are stable after exposure to ultrasound, high ionic strength and extreme pH conditions, and revealed the presence of basic groups associated to the AuNPs surface. Electrophoretic and dynamic light scattering indicated that biogenic dispersions of AuNPs are stabilized by a steric mechanism. The AuNPs produced by *C. metallidurans* CH34 are not cytotoxic towards bacterial cells, in contrast to biogenic AgNPs. These stable non-toxic biogenic AuNPs have potential clinical applications including development of topic delivery formulations and optical biosensors.

## 1 Introduction

The need for cost-effective eco-friendly synthesis of metallic nanoparticles has established new methods to replace the chemical synthesis. The biogenic synthesis of nanoparticles is of increasing interest. Bacteria are versatile biocatalysts for detoxification and biotransformation of heavy metals and aromatic compounds [1–8]. Microbial systems have been selected for synthesis of nanoparticles due to their detoxification mechanisms of metallic ions through extracellular or intracellular reduction. Biogenic synthesis of metallic nanoparticles by diverse microorganisms has been described [9, 10]. Accordingly, synthesis of gold nanoparticles by *Pseudomonas, Shewanella* and *Streptomyces* strains has been reported [11–15].

*C. metallidurans* is a model heavy metal-resistant bacterium that harbours gene clusters enabling detoxification of diverse heavy metal ions and complexes [1, 2, 16, 17]. Interestingly, *C. metallidurans* is also involved in the biogeochemical cycle of gold. Presence of *C. metallidurans* cells in biofilms that covered the surface of gold grains has been reported [18, 19]. Transcriptomic analysis suggested that oxidative stress and heavy metal resistances genes including an Au-specific operon were involved in Au reduction by *C. metallidurans* [19]. Recently, formation of gold nanoparticle and microparticle aggregates onto the surface *C. metallidurans* biofilms has been reported [20].

The aim of this study was to obtain a stable dispersion of AuNPs by *C. metallidurans* CH34 cells. Bacterial cultures were fractionated and exposed to a gold aqueous solution. Using this procedure extracellular dispersions of NPs were obtained. The biogenic colloid of dispersed AuNPs was characterized by surface plasmon spectroscopy, dynamic and electrophoretic light scattering and nanoparticle tracking analysis. Solid state AuNPs were characterized by powder X-ray diffraction, Fourier transform infrared spectroscopy and TEM. Stability of the colloid was characterized by electrophoretic and dynamic light scattering. A synthesis mechanism is proposed. Finally, cytotoxic effect of the biogenic gold NPs on bacterial growth was evaluated.

## 2 Experimental

### 2.1 Materials

Solid HAuCl_4_·3 H_2_O (99.9% purity) was obtained from Sigma Aldrich (Saint Louis, MO, USA). NaCl, NaOH and KCl were obtained from Merck (Darmstadt, Germany). Beef extract and yeast extract were acquired from Becton Dickinson (Cockeysville, MD, USA) and agar solid medium was obtained from Merck (Darmstadt, Germany). Biogenic AgNPs (40 nm average diameter) were recovered from fungal *Fusarium oxysporum* filtrates [13].

### 2.2 Biosynthesis of gold nanoparticles

*C. metallidurans* CH34 [21] and *E. coli* MG1655 [22] were grown in a low ionic strength medium with meat extract (20 g L^-1^) and yeast extract (20 g L^-1^) [23]. Biogenic synthesis of gold nanoparticles was done according to methods A and B [13], with modifications (Fig. 1S). Method A evaluates the effect of bacterial biomass on the biogenic synthesis of AuNPs, and method B evaluates the effect of bacterial extracellular medium on the gold reduction process. In method A, biomass (40 mL) was directly exposed to a gold solution achieving a final concentration of 2 mM. In method B, bacterial biomass was used to obtain bacterial filtrates that were exposed to a gold solution (2 mM). In both methods, cell cultures grown until stationary phase were used as starting bacterial biomass. For method A, cells were collected by centrifugation and washed 3 times with doubly deionized water. Cell pellet was suspended in doubly deionized water and diluted up to a concentration of 3.3 × 10^9^ CFU mL^-1^. The bacterial suspension was exposed to AuCl_4_^−^ (2 mM) and then incubated during 96 h in darkness without shaking at 30° C. The HAuCl_4_ stock solution (40 mM) was previously adjusted to pH 7 with NaOH. To recover the biogenic colloid, the bacterial suspension was centrifuged at 5,000 × *g* during 10 min and supernatant was filtered using a 0.45 μm pore size nitrocellulose membrane. The dispersion of biogenic AuNPs was recovered in the permeate fraction. For method B, cells from a stationary phase bacterial culture were collected by centrifugation and washed 3 times with doubly deionized water. Cell pellet was suspended in doubly deionized water and diluted up to a concentration of 6 × 10^9^ CFU mL^-1^. To obtain the bacterial secretome, biomass was incubated during 3 days in flasks with shaking (150 rpm) at 30° C. Cells were centrifuged at 5,000 × *g* during 10 min and the supernatant was recovered and filtered using a 0.2 μm pore size nitrocellulose membrane. The bacterial secretomes were recovered [24] and incubated with AuCl_4_^−^ (2 mM) during 96 h in darkness without shaking at 30° C.

**Fig. 1.**
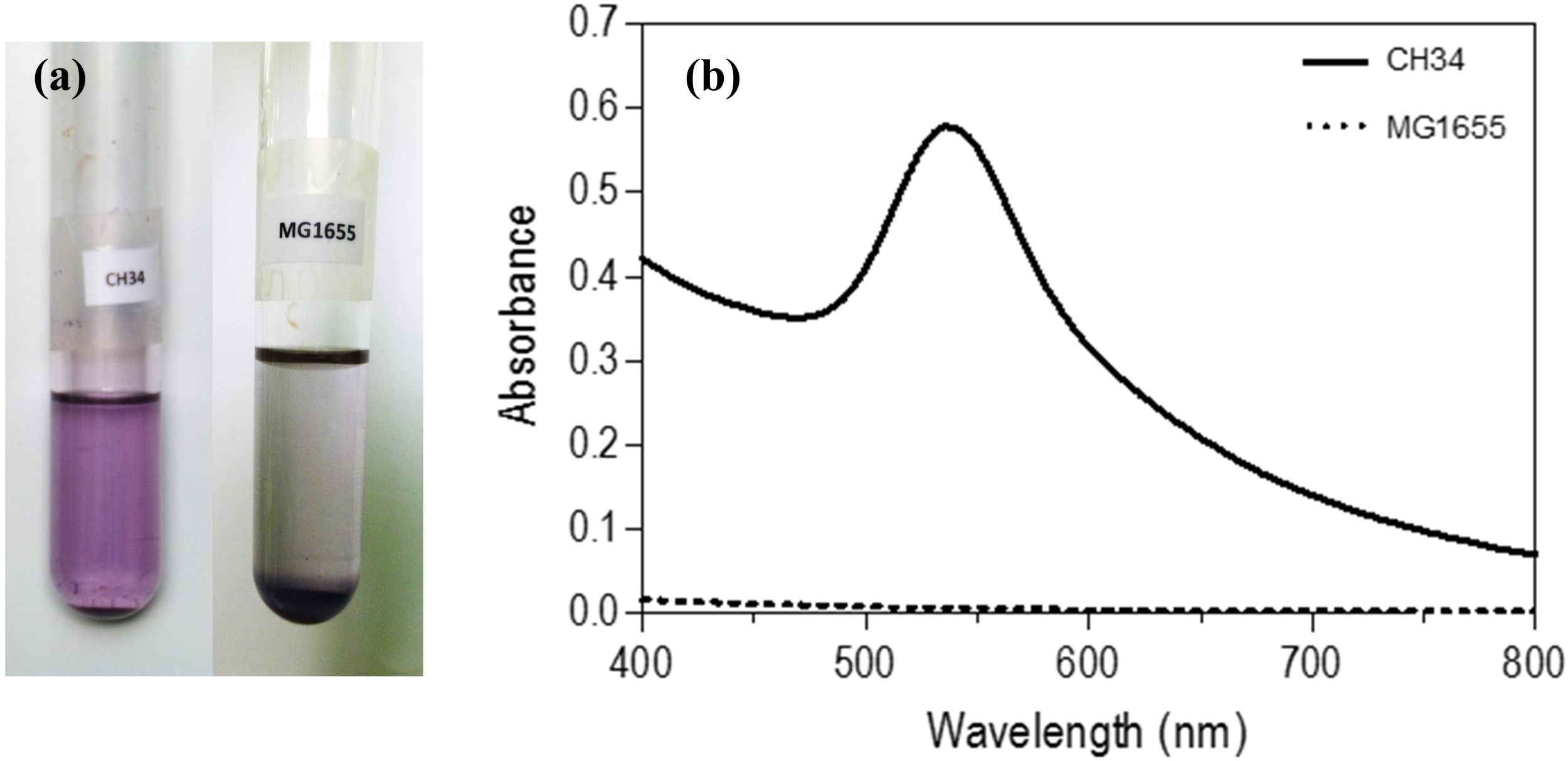
Optical properties of gold nanoparticles synthetized by bacteria. (a) Reduction of AuCl_4_^−^ by *C. metallidurans* CH34 (left) and *E. coli* MG1655 cells (right). (b) Visible spectra of the aqueous fractions obtained from *C. metallidurans* CH34 and *E. coli* MG1655 supernatants.

### 2.3 Characterization of biogenic gold nanoparticles

The UV-visible absorbance spectra of gold nanoparticles were obtained at room temperature with an Agilent 8453 spectrophotometer equipped with a diode array (Palo Alto, CA, USA). Hydrodynamic diameter (Z-Ave) and dispersion index (PdI) of the biogenic dispersions were determined by dynamic light scattering (DLS) measurements, and zeta-potentials (ζ-potentials) were measured by electrophoretic light scattering using a ZetaSizer Nano ZS instrument (Malvern, Worcestershire, UK). Additional particles size distributions and concentration were determined by nanoparticle tracking analysis (NTA) technology using a NanoSight LM20 device (Amesbury, UK). In all cases, temperature was fixed at 25° C and 1 mM KCl solution was used as dissolvent. Morphology and size distribution of the biogenic AuNPs was observed by transmission electron microscopy. Images were obtained using a Carl Zeiss (Libra) transmission electron microscope (TEM) operating at 120 keV and analyzed using software ImageJ 1.47v. To obtain the images, one drop of biogenic AuNPs was deposited on a carbon-coated parlodion film supported in 300 mesh copper grids (Ted Pella, Redding, CA, USA). For EDX measurements, biogenic AuNPs were extensively washed with MilliQ water, concentrated by centrifugation, and deposited on a carbon grid. Spectra were acquired with a Carl Zeiss, EVO MA-10 device. Solid state characterization was done to powder samples of the biogenic AuNPs. To obtain the powder samples, colloidal dispersions of biogenic AuNPs were centrifuged at 20,000 × *g* during 20 min and then washed 3 times with MilliQ water. The resulting sediment containing the biogenic AuNPs was lyophilized and then analyzed by powder X-ray diffraction (XRD) and Fourier transform infrared (FT-IR) spectrometry. Powder XRD spectra were obtained in a Shimadzu XRD 6000 diffractometer using Cu Kα radiation (1.5406 Å) operating at 30 mA and 40 k V. Scan speed was 0.02 degree min^-1^ and time constant was 2 s. FT-IR spectra of powder samples were recorded using a Bomem MB spectrometer in the 4,000-400 cm^-1^ frequency range using an attenuated total reflectance mode. A total of 250 scans and a resolution of 4 cm^-1^ were employed to obtain each spectrum. Concentrations of gold and silver in the nanoparticles dispersions were determined by inductively coupled plasma (ICP) Perkin Elmer Optima 3000 DV spectrometry.

### 2.4 Cytotoxic activity assays

The cytotoxic effect of biogenic AuNPs and Ag nanoparticles on *E. coli* MG1655 cells was evaluated by minimal inhibition concentration (MIC) measurements. MIC of biogenic nanoparticles was determined by the microdilution method [25]. The liquid medium used in these assays contained meat extract (5 g L^-1^) and yeast extract (5 g L^-1^) and was inoculated to a final bacterial concentration of 5 × 10^5^ CFU mL^-1^. Aliquots of bacterial cultures were exposed to equivalent concentrations of gold and silver NPs obtained by serial dilutions of the colloids. The initial concentration of both Au and Ag nanoparticles was 14.5 μg mL^-1^. MIC was defined as the minimum concentration of nanoparticles that inhibit the bacterial growth in liquid medium after an incubation period of 16 h at 37° C. The effect of gold and silver nanoparticles on bacterial growth was visually analyzed on agar plates. Solid medium contained meat extract (5 g L^-1^), yeast extract (5 g L^-1^) and agar (15 g L^-1^). Agar plates (85 mm diameter) were seeded with an aliquot (100 μL) of 1 × 10^8^ CFU mL^-1^ of MG1655 cells and exposed to AuNPs and AgNPs. The nanoparticle dispersions were serially diluted and drops containing 20 μL of each dilution were poured onto the previously seeded solid medium. Agar plates were incubated during 16 h at 37° C.

## 3 Results

### 3.1 Biogenic synthesis of gold nanoparticles

After 96 h in darkness, supernatant of strain CH34 cell culture developed a purple color indicating the production of an extracellular dispersion of biogenic AuNPs (Fig. 1a). No viable cells were detectable after the biosynthesis process, whilst cell viability of CH34 cultures without Au(III) ions remained constant. In contrast to *C. metallidurans* CH34, development of a purple color in supernatant of *E. coli* MG1655 cells was not detected and accumulation of a dark sediment was observed (Fig. 1a). Viable *E. coli* cells were not detected after incubation with Au(III) ions, whereas cell viability of control cultures without Au(III) ions remained constant.

The reduction of Au(III) ions and production of biogenic AuNPs by the secretomes of strains CH34 and MG1655 was studied. Incubation of both bacterial secretomes with Au(III) ions did not developed a purple color and appearance of an insoluble yellow gold complex was observed (not shown).

### 3.2 Characterization of biogenic AuNPs in colloidal state

The visible spectrum of supernatants of CH34 cultures obtained after incubation with Au(III) ions showed a maximum absorbance peak at a wavelength of 536 nm (Fig. 1b). No absorbance was detected in supernatants of *E. coli* MG1655 cells (Fig. 1b) and secretomes of both strains. These results indicate that under these conditions only CH34 cells are capable to produce extracellular dispersions of biogenic AuNPs.

The biogenic colloid obtained with CH34 cells was characterized by light scattering measurements. Dynamic and electrophoretic light scattering measurements indicated that the AuNPs dispersion presents a Z-Ave of 126.4 nm (Fig. 2a) with an associated zeta-potential of -0.25±3 mV. Additional measurements using NTA technology correlated well with previous DLS results. Mean diameter of the biogenic colloid was 124 nm, with a mode of 78 nm and the presence of minor populations of 302 nm and 405 nm (Fig. 2b, video in Supplemental information). Concentration of biogenic AuNPs diluted 4 times with 1 mM KCl and reported by NTA was 4.19 × 10^9^ particles mL^-1^. Therefore, the biogenic colloid has a concentration of 1.68 × 10^9^ particles mL^-1^.

**Fig. 2.**
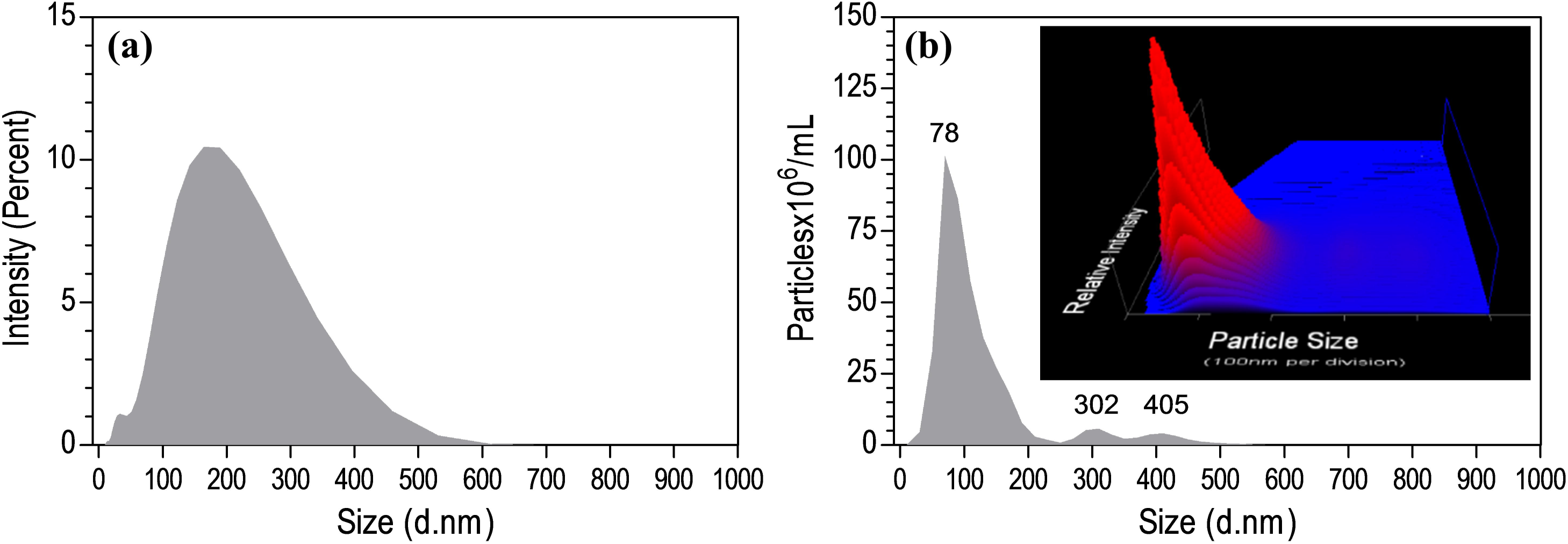
Comparison of size distributions of gold nanoparticles synthetized by *C. metallidurans* CH34 obtained by light scattering measurements. (a) Size distribution obtained by dynamic light scattering (DLS) measurements. Reported Z-Ave of 126.4 nm and PdI of 0.284 (b) Size distribution and particles concentration obtained by nanoparticle tracking analysis (NTA). Reported mean of 124 nm, mode of 78 nm and SD of 88 nm.

Stability of the biogenic AuNPs colloid after exposure to ultrasound, extreme pH conditions and elevated ionic strength was evaluated by DLS measurements (Table 1). Hydrodynamic diameter of control samples was 126.4 nm with associated PdI of 0.284. Ultrasound treatment did not affect size distribution the colloid, with associated Z-Ave and PdI values of 124.9 nm and 0.279, respectively. Stability of the colloid was also evaluated in the presence of highly acidic (pH 1) and alkaline (pH 10) conditions. The presence of an acidic environment reduced Z-Ave from 126.4 nm to 114.6 nm, whilst an alkaline environment increased Z-Ave to 135.3 nm. In both cases, PdI of the colloid increased to 0.33. Finally, presence of elevated ionic strength (500 mM NaCl) showed no-effect on size distribution, with associated Z-Ave and PdI values of 127.5 nm and 0.286, respectively.

**Table 1.** Dynamic light scattering measurements of biogenic AuNPs dispersions produced by *C. metallidurans* CH34 after exposure to ultrasound, extreme pH and elevated ionic strength (average of 3 measurements)

| Experimental condition | Z-Ave (d.nm) | PdI |
| --- | --- | --- |
| Control | 126.4 | 0.284 |
| Ultrasound | 124.9 | 0.279 |
| pH 1 | 114.6 | 0.332 |
| pH 10 | 135.3 | 0.334 |
| NaCl (500 mM) | 127.5 | 0.286 |

### 3.3 Characterization of biogenic AuNPs in solid state

The degree of crystallinity of the biogenic gold nanoparticles was determined by powder X-ray diffraction measurements. XRD spectrum showed Bragg´s reflection peaks at positions 2θ = 38.1°, 44.3°, 64.6°, 77.5° and 81.7° (Fig. 3a). Position of the reflection peaks can be associated to diffraction planes (111), (200), (220), (311) and (222) of the crystalline unit cell of elemental gold as seen in the 04-0784 file of the JCPDS database. The all odd or all even Miller index values of the diffraction planes indicate that the biogenic gold nanoparticles have a face-centered cubic (*fcc*) crystalline structure. The calculated lattice constant (a) using the distance value (d) associated to the {220} diffraction planes was estimated to be 4.1 Å. This value agrees with standard report value from JCPDS (a = 4.1 Å), confirming that biogenic AuNPs synthetized by strain CH34 are composed of elemental gold. Predominant diffraction plane present in the sample was also determined. Ratio of the intensity peaks (200), (220), (311) and (222) with respect to (111) are lower than ratio of the intensity reported in the standard JCPDS gold sample (0.25 versus 0.52, 0.18 versus 0.32, 0.14 versus 0.36, and 0.04 versus 0.12, respectively). This analysis indicates that the crystallographic structure of AuNPs synthetized by strain CH34 is dominated by {111} diffraction planes.

**Fig. 3.**
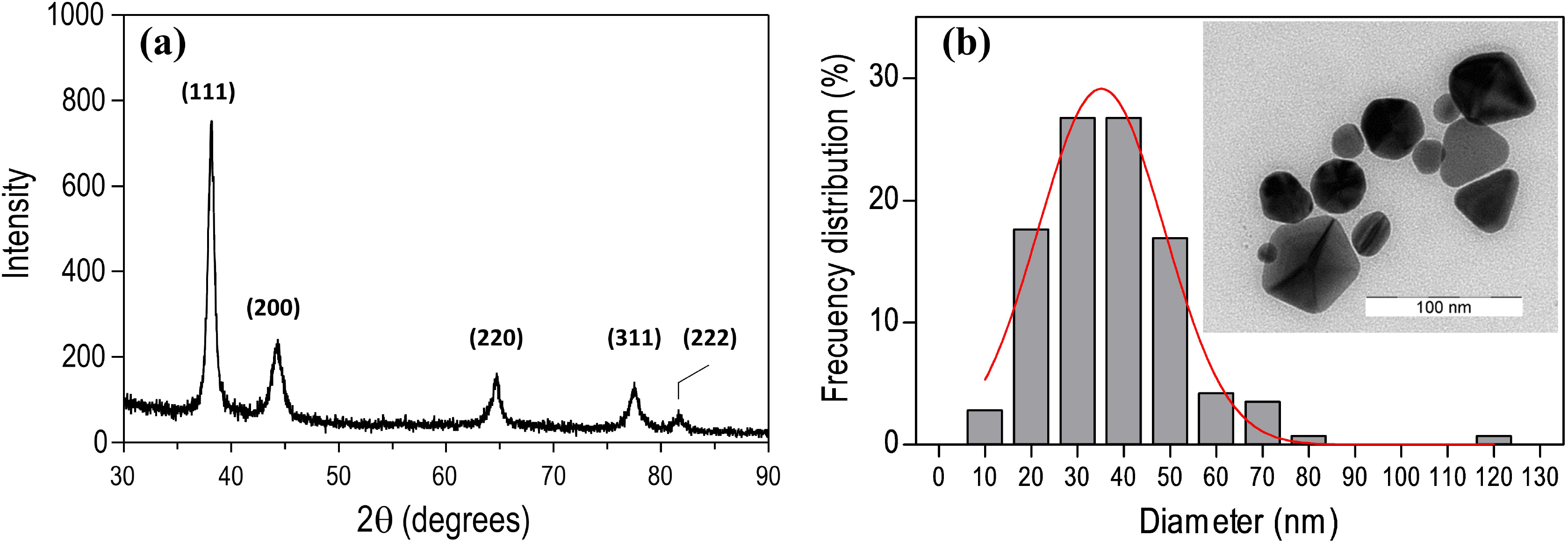
Physical properties of biogenic gold nanoparticles synthetized by *C. metallidurans* CH34. (a) Powder X-ray diffraction spectrum. (b) Size frequency distribution and morphology of the AuNPs determined by analysis of TEM images (n=142). Mean of 37.1 with SD of 15.5 nm. Inset: representative field of biogenic AuNPs.

The morphology and size distribution of the biogenic AuNPs were studied by transmission electron microscopy. Morphology of the biogenic AuNPs was dominated by decahedral and triangular nanostructures (Fig. 3b). The metallic core of biogenic AuNPs showed diameters ≥ 10 nm, with 85% of the particles showing a diameter between 20 and 60 nm. Few truncated triangular nanoplates with a diameter ≥ 70 nm were also observed. Mean of the frequency distribution was 37.1±15.5 nm. Further analysis of TEM images showed that perimeter of all nanoparticles was surrounded by a layer of non-metallic material with a width of ~ 4.5 nm (Fig. 3b (Inset)). These structures could be organic capping ligands adsorbed on the surface of the biogenic AuNPs.

Energy-dispersive X-ray analysis of biogenic AuNPs showed the presence of sulfur atoms associated to AuNPs (Fig. 4a). Afterwards, the chemical nature of the biogenic AuNPs was characterized by FT-IR spectroscopy (Fig. 4b). Absorbance spectrum of the biogenic AuNPs showed several peaks that correlate with molecular vibrations of functional groups from proteins; therefore, biogenic AuNPs spectrum was analyzed based on the FT-IR spectrum of bovine serum albumin [26]. The broad band between 3700 and 3000 cm^-1^ corresponds to vibrations of water molecules and protein functional groups. The higher peak at 3450 cm^-1^ corresponds to OH··· stretching vibrations of residual H_2_O (*v*_O-H_…). The following absorption bands at lower energy correspond with amide A group of vibrations. These are the stretching vibrational modes of N-H group of the peptide bonds (*v*_N-H_…; 3290 cm^-1^ and *v*_N-H_; 3100 cm^-1^). The absorption band between 3000 and 2800 cm^-1^ corresponds with stretching vibrations of C-H groups. The broad band between 1600 and 1700 cm^-1^ corresponds with molecular vibrations of amide I group. This group includes stretching modes of hydrogen bond acceptor (*v*_C=O_…; 1675, 1665 and 1630 cm^-1^) and hydrogen bond non-acceptor (*v*_C=O_; 1695 cm^-1^) molecular vibrations associated to C=O group of peptide bonds. The absorption band between 1600 and 1500 cm^-1^ corresponds with molecular vibrations of amide II group. This group includes bending modes of the hydrogen bond acceptor (δ_N-H_…; 1550 and 1520 cm^-1^) and hydrogen bond non acceptor (δ_N-H_; ≈ 1500 cm^-1^) molecular vibrations associated to N-H group of peptide bonds. A similar absorbance pattern has been described when albumin is adsorbed onto chemically synthetized AuNPs [27, 28]. The absorption band between 1400 and 1360 cm^-1^ corresponds with symmetric stretch (*v*^S^_COO^-^_) of the carboxylate groups of Asp and Glu residues. This global data suggests that dispersions of biogenic AuNPs synthetized by strain CH34 are covered by a layer of proteins.

**Fig. 4.**
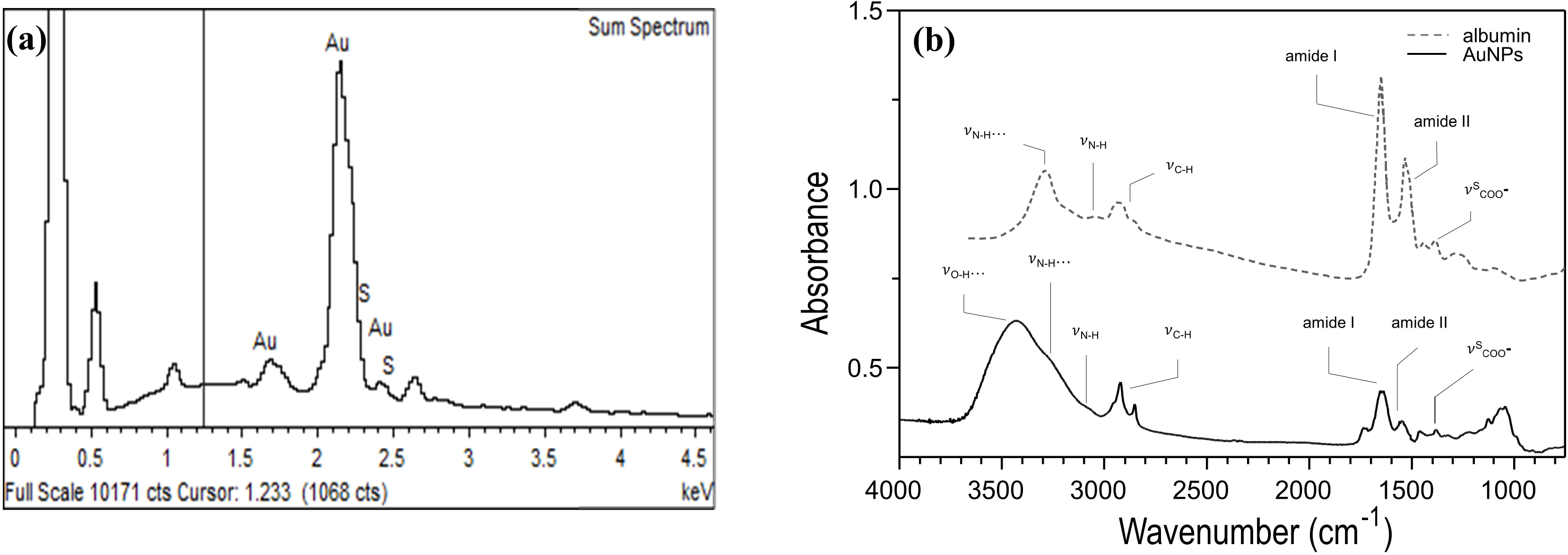
Chemical characterization of biogenic AuNPs synthetized by *C. metallidurans* CH34. (a) EDX spectrum of biogenic AuNPs. (b) FT-IR spectrum of biogenic nanoparticles (straight line), and bovine serum albumin adapted from [26] (dotted line).

### 3.4 Effect of biogenic AuNPs on bacterial growth

The effect of biogenic gold and silver NPs on bacterial cell growth was compared. Microdilution method indicated that biogenic AuNPs did not affect the growth of *E. coli* MG1655 cultures, while biogenic AgNPs showed a MIC of 0.45 μg mL^-1^ (Fig. 5). This value is in agreement with previous reports [10, 29, 30]. Similar results were observed with biogenic Au and Ag nanoparticles on solid growth medium. These results indicate that biogenic AuNPs synthetized by *C. metallidurans* strain CH34 do not affect growth of *E. coli* MG1655. Therefore, these biogenic AuNPs are not cytotoxic for *E. coli* cells.

**Fig. 5.**
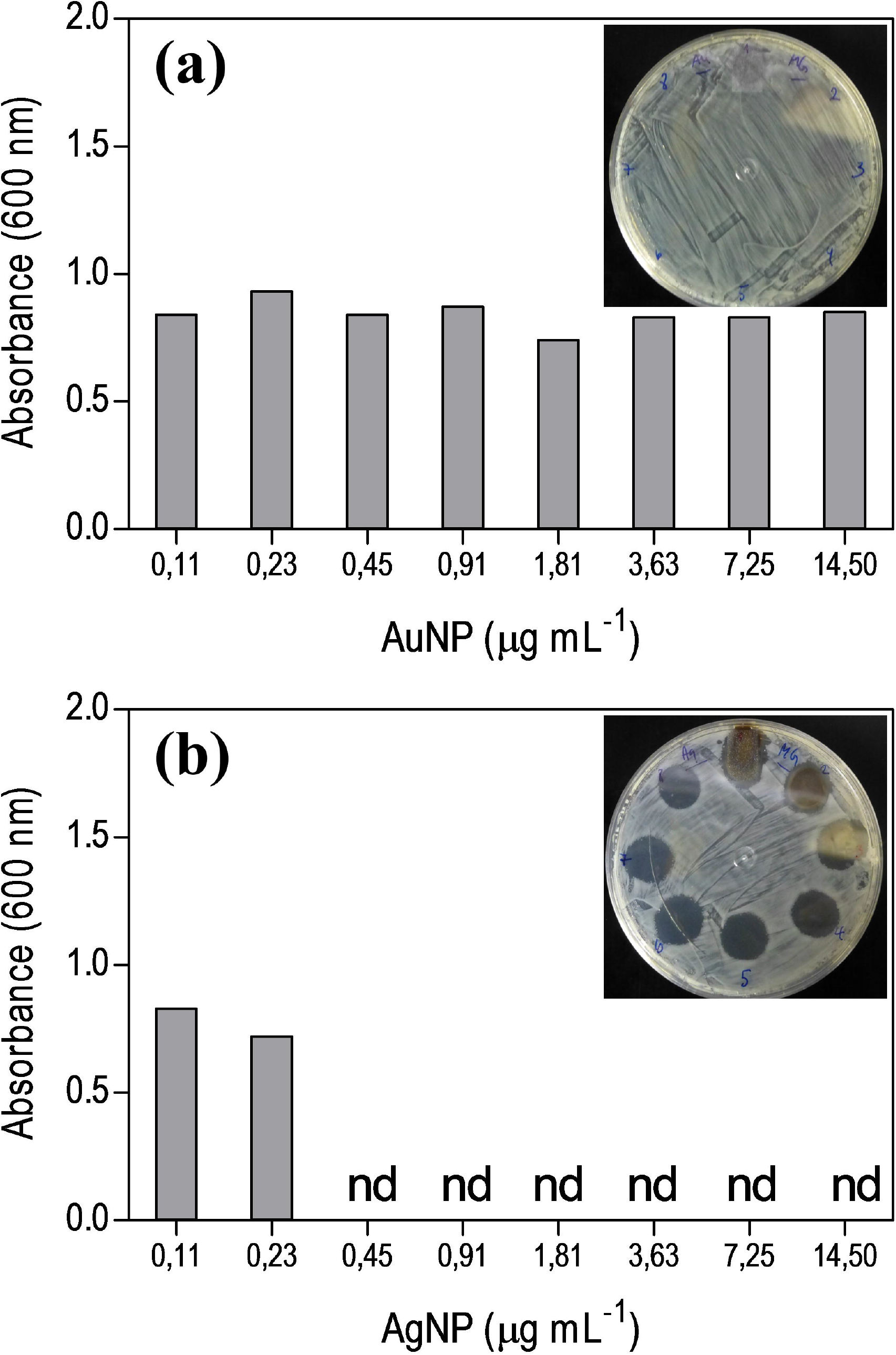
Minimal inhibitory concentration of AuNPs on planktonic cultures of *E. coli* MG1655. a, AuNPs produced by *C. metallidurans* CH34 b, AgNPs produced by *Fusarium oxysporum*. nd: no growth detected. Inset: growth of *E. coli* MG1655 cells on agar plates after exposure to an equivalent amount of NPs.

## 4 Discussion

In this study, stable dispersions of AuNPs were produced after reduction of Au(III) ions by *C. metallidurans* CH34 cells. Production of biogenic metallic nanoparticles occurred by redox agents allocated in the cellular biomass. Exposure of *C. metallidurans* CH34 and *E. coli* MG1655 secretomes to Au(III) ions did not produce AuNPs dispersions. Therefore, under these experimental conditions both strains do not secrete enzymes or compounds to the extracellular medium that allow reduction of Au(III) ions with concomitant generation of biogenic AuNPs dispersion. Reduction of Au(III) ions by *E. coli* cells has been reported, but AuNPs aggregate onto bacterial cells and do not generate extracellular dispersions [31]. Production of extracellular AuNPs by cyanobacterium *Plectonema boryanum* UTEX485 cells has been described [32].

These results indicate that the mechanism of Au(III) reduction during the biogenic synthesis of extracellular dispersions of AuNPs implies oxidation of molecules that are produced by strain CH34 and are absent in strain MG1655. It is propose that reduction of Au(III) ions and production of extracellular dispersions of AuNPs is mediated by oxidation of macromolecules that are located in the cellular membranes or periplasm of CH34 cells and are absent in MG1655 cells. Accordingly, a comparative bioinformatic analysis of both genomes indicated that, in contrast to strain MG1655, *C. metallidurans* CH34 possess a *cop* gene cluster located within the megaplasmid pMOL30 that encodes periplasmic and outer membrane sulfur rich proteins that contain a significant number of methionine (Met) and cysteine (Cys) residues (CopA 36 Met/1 Cys, CopB 51 Met, CopC 7 Met, CopK 9 Met, CopJ 4 Met/2 Cys) (Fig. 6). Moreover, proteomic studies of strain CH34 demonstrated the constitutive expression of the periplasmic protein CopK and the outer membrane bound protein CopB [33], showing that these proteins are highly expressed in cells of *C. metallidurans* CH34.

**Fig. 6.**
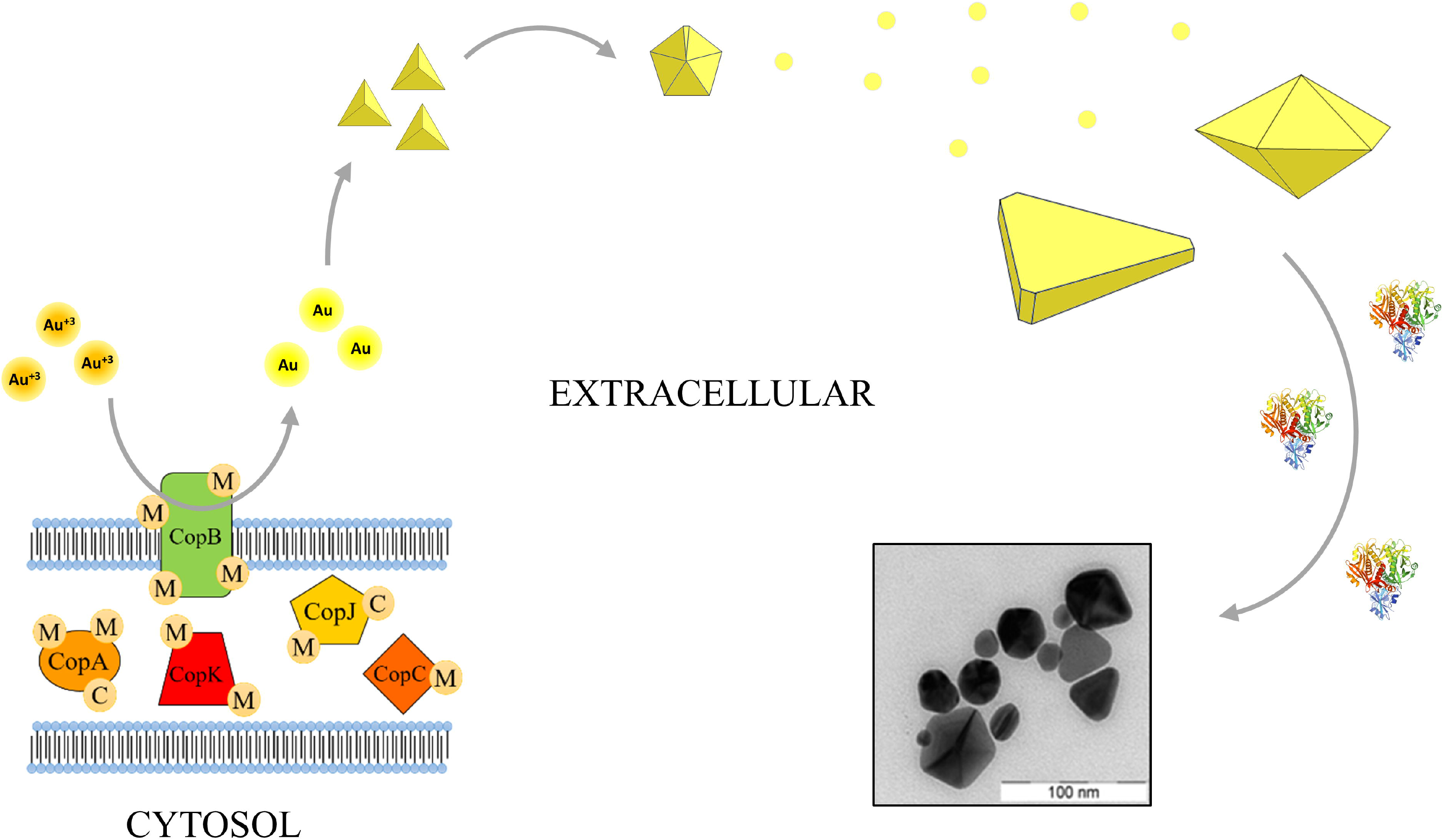
Proposed mechanism for the reduction of Au(III) ions by *C. metallidurans* CH34 cells and subsequent formation of extracellular dispersions of AuNPs.

Thus, during biogenic synthesis experiments, the presence of Cop sulfur rich proteins located at the membranes and periplasm of CH34 cells might allow direct interaction between the sulfur atoms of methionine and cysteine residues and the diffusive Au(III) ions present in the extracellular space. In this manner, it is proposed that extracellular Au(III) ions oxidize the sulfur atom of the methionine residues into methionine-sulfoxide. This event has been described for methionine residues from ribonuclease A and glycyl-D,L-methionine dipeptide, with concomitant production of Au(0) [34–36]. In addition, Au(III) ions may oxidize cysteine to cystine, and subsequently to sulfonic acid [37, 38]. After interaction, the reduced gold atoms might return to the medium and generate nucleation centers that finally evolve into extracellular AuNPs dispersions (Fig. 6). Moreover, under physiological conditions, redox cycling enzymes such as Met sulphoxide reductases might reduce and recover the functionality of oxidized sulphur containing residues. Therefore, this analysis suggests that Met residues of CopA and CopB are especially involved in the reduction of gold ions to produce biogenic extracellular AuNPs; and that a redox cycling process of these residues, probably mediated by Met sulphoxide reductases, is involved in the biochemical synthesis of gold nuggets in nature [19].

Analysis of electron microscopy images showed that nanoparticles morphology was dominated by decahedral and triangular structures. Origin of these structures may occur by interaction of ultra-small tetrahedral nanoparticles [39-41]. Tetrahedral morphology becomes unstable when nanoparticles reach diameters close to 10 nm. Then, interaction of five tetrahedral units lowers down the surface area allowing formation of decahedral structures [39] (Fig. 3b and Fig. 6). On the other hand, presence of major triangular nanoplates has been attributed to a slow kinetic process during nanoparticles formation [42]. After long periods, interaction of decahedra like structures serve as nucleation centers that finally evolve into triangular and truncated triangular structures. It is propose that the synthesis mechanism of biogenic AuNPs involves an initial step where small tetrahedral nanoparticles are generated. The interaction of small tetrahedral units evolves into decahedral structures. As observed by TEM, one population of these decahedral structures may serve as nucleation centers for the subsequent biosynthesis of triangular nanoplates and other population might evolve up to reach diameters close to 50 nm.

Analysis of XRD spectrum showed a bias towards {111} diffraction planes. This non-ideal behavior may be attributed to morphological properties of decahedral and triangular nanoparticles observed by TEM. Decahedral structures are formed by interaction of five tetrahedral particles bounded by its lower energy {111} twin planes [39, 41]. Also the major faces of triangular and triangular like nanoplates present a {111} crystallite structure [40, 43, 44]. As the surface area of these nanostrucutres is dominated by {111} diffraction planes, it is propose that morphology of the biogenic AuNPs contains the information that enhanced the signal of (111) diffraction plane, leading to a XRD spectrum dominated by {111} diffraction planes as described above.

Comparison of size distribution of the biogenic AuNPs obtained by light scattering measurements using two different technologies showed similar results. Dynamic light scattering measurements indicated that Z-Ave was 126.4 nm, whilst NTA indicated that mean diameter of the distribution was 124 nm. These results do not correlate with size distribution deduced from TEM images (average diameter of 37.1 nm). The difference between diameters determined by TEM and light scattering technologies can be explained by the characteristics of each technique. Size distributions obtained from TEM images are proportional to the length of the nanoparticles and do not include the width of capping ligands adsorbed onto AuNPs surface. Also, size distributions obtained from TEM images reflect the results from the analysis of AuNPs samples that are in a solid static state. In contrast, size distributions obtained from DLS and NTA measurements are proportional to the volume of the nanoparticles and represent the accumulation of multiple measurements obtained from a nanoparticles population in dynamic state. Size distributions obtained by these light scattering technologies reveal intermolecular interactions between AuNPs dispersions that leads to multiple equilibriums and are finally detected as an average of major sized nanoparticles complexes [28, 45-47]. Therefore, size distributions of colloids obtained by DLS and NTA contain information about interacting and non-interacting nanoparticles, and detect small amounts of larger nanoparticles that are outside the normal distribution and are not captured in TEM images [48].

The stability of the biogenic AuNPs colloid was determined by exposure to ultrasound, extreme pH conditions and elevated ionic strength by DLS measurements. Ultrasound fuses non-functionalized AuNPs into worm like units with concomitant increment in Z-Ave and PdI [49]. Exposure to prolonged sonication periods did not change Z-Ave nor PdI values, indicating that the biogenic colloid is not sensitive to ultrasound perturbation by the protective effect of a layer of capping ligands that cover the surface of the particles. Further Z-Ave determinations after exposure to extreme pH conditions indicated that the biogenic colloid endures the chemical perturbance of acidic and alkaline environments and reveals the presence of basic functional groups located at the surface of the nanoparticles. Exposure to an acidic environment reduced the hydrodynamic diameter and increased PdI of the colloid. It is propose that low pH conditions generate a re-distribution of the interacting nanoparticles population. Protonation of basic groups of the capping ligands increase electrostatic repulsion of at least one fraction of the interacting nanoparticles. This process probably generates a novel group of non-interacting nanoparticles that splits the population distribution and is detected as a reduction in Z-Ave and an increment in the PdI of the colloid. On the other hand, alkaline environment increased both Z-Ave and PdI of the colloid. Deprotonation of basic groups might reduce the electrostatic repulsion of at least one fraction of the interacting nanoparticles. This process would generate a group of highly-interacting nanoparticles that splits the former population distribution and is detected as an increment in both Z-Ave and PdI of the biogenic colloid. The presence of high concentrations of NaCl did not alter size distribution of the colloid. This result demonstrates that biogenic AuNPs do not flocculate nor aggregate in the presence elevated ionic strength conditions, and indicates that biogenic AuNPs colloids obtained with *C. metallidurans* CH34 cells would remain stable even under physiological conditions (150 mM NaCl).

The stability mechanism of the biogenic colloid was studied by electrophoretic and DLS measurements in the presence and absence of ionic strength. Elevated absolute ζ-potential values (≥ 30 mV) are indicative of electrostatic repulsion between nanoparticles and therefore to the stability of the colloid mediated by an electrostatic repulsion mechanism [50]. In this case, presence of ionic strength shields the electrostatic repulsion between the particles, and induces flocculation and increment in Z-Ave of the colloid. The ζ-potential of biogenic AuNPs synthetized by strain CH34 was ~ 0 mV (-0.25±3 mV), and Z-Ave did not change in presence of 500 mM NaCl (Table 1). These results indicate that the biogenic colloid is not stabilized by electrostatic repulsion forces and suggests that the stability mechanism is based on a steric repulsion between the capping ligands of the nanoparticles. Stabilization by steric repulsion mechanism indicates that elevated molecular weight ligands are adsorbed onto the surface of the biogenic AuNPs [28, 51], and supports FT-IR results that reveal the presence of functional groups of proteins.

## 5 Conclusions

Stable colloidal dispersions of AuNPs were obtained after incubation of *C. metallidurans* CH34 cells with Au(III). Potential CH34 proteins such as CopaA, CopB or CopK may act as electron donors for the reduction of Au(III) ions during the biosynthesis process, and a mechanism for the production of extracellular AuNPs by strain CH34 was proposed.

Average diameter of the biogenic AuNPs in colloidal state obtained from light scattering measurements (DLS and NTA) was four times higher than diameter of the biogenic AuNPs in solid state obtained from analysis of TEM images. This comparison shows that the size distributions obtained from analyses of nanoparticles in colloidal and solid state are not comparable.

The biogenic colloid is stable under chemical and physical perturbations, which is useful for potential applications. Clinical applications include the topic delivery of bioactive compounds that may interact with the protein layer of the nanoparticles and the development of colorimetric biosensors that require colloidal probes with elevated stability and absorptivity.

---

**Fig. 1S.**
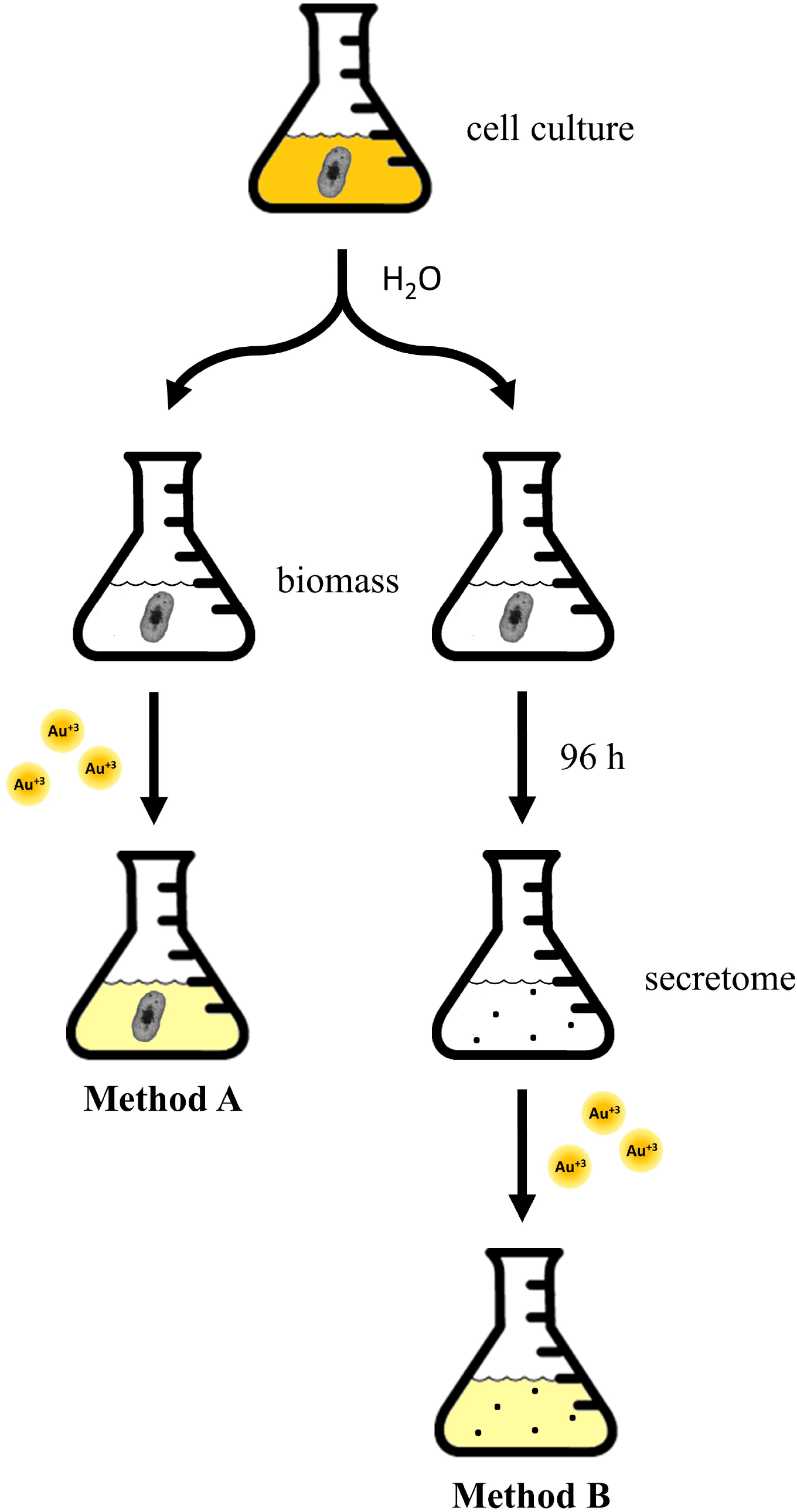
Workflow of cell culture fractionation. **Interactive media S1. Video showing light dispersion and Brownian motion of biogenic AuNPs synthetized by *C. metallidurans* CH34 cells**.

